# Visual impairment, spectacle wear and academic performance among junior high school students in western China

**DOI:** 10.1101/2019.12.13.875153

**Authors:** Jingchun Nie, Wei Nie, Ming Mu, Lifang Zhang, Shuyi Song, Lanxi Peng, Qiufeng Gao, Jie Yang

## Abstract

In September 2013, 2248 students from 36 junior high schools in Shaanxi Province underwent assessment of visual acuity (VA), completed a questionnaire about their spectacle use and were administered a standardized mathematics examination. Among 2,248 students (mean age 13.6 years, 52% male), visual impairment was present in 699 (31%, 95% Confidence Interval 29-33%). Spectacle wear was observed in 360 of 966 children needing glasses (37%). Ownership of spectacles among children needing glasses was associated with poorer uncorrected vision in the better-seeing eye (P <0.001) and paternal educational (p=0.001), but not age, sex, boarding at school, both parents having out-migrated for work or maternal education. Spectacle ownership among children with visual impairment was associated with better test performance (P=0.035). Therefore, visual impairment and non-wear of spectacle were common. Wearing spectacles was associated with better academic performance in this cross-sectional analysis, consistent with recent trial results among younger children.

## Introduction

In both rural and urban areas of China, the prevalence of visual impairment due to refractive error among children is high,^1,2^ and rises with age.^3^ Despite the ease and safety of correcting refractive error with prescription eyeglasses, a significant proportion of rural children needing spectacles in China neither own nor wear them.^4–7^

Many previous reports on refractive error prevalence among Chinese children are either rely on samples from limited areas or employ unclear sampling methodology.^5^ A small number of studies have sampled western rural areas, but have focused on primary schools students.^5,8^ To our knowledge there has not been a large, population-based study of visual impairment among secondary school students in western China, home to approximately 15 million 13 to 15 year-olds.^9^

Uncorrected refractive error has been associated with reversible decrement in self-reported visual function among school-aged children.^10^ In addition, two recent randomized, controlled trial have demonstrated statistically significant improvement in academic performance with provision of free spectacles.^11,12^ However, data linking spectacle ownership/wear and school performance remain sparse, particularly in the large populations of Asia, where rates of uncorrected refractive error are high.^13–15^ Factors determining the purchase and wear of spectacles are also not well understood.

We report on the results of a study of visual impairment, spectacle wear and academic performance in a randomly sampled population of junior high school students in Shaanxi, a middle-income province^9^ in western China.

## Methods

The protocol for this study was approved in full by Institutional Review Boards at Stanford University (Palo Alto, USA, No.24847). Permission was received from the local Board of Education and the principals of all schools involved. The principles of the Declaration of Helsinki were followed throughout.

### Sampling

In September-October 2013, a list of all 36 rural junior high schools in three randomly selected counties (Yuyang, Hengshan and Jingbian) in Yulin prefecture was obtained, and all children in one randomly-selected class in each of the 7^th^ and 8^th^ grades were enumerated.

### Questionnaires

Enumerators (trained graduate students from Shaanxi Normal University) administered a questionnaire to collect information on student and family characteristics, including grade level, sex, boarding status at school, and parental schooling and migration status. The same team also administered a timed (25 minutes) and proctored mathematics examination to all students in the selected classes. Items were selected with the help of local educators from a bank of questions developed by the Trends in International Mathematics and Science Study (TIMSS) testing service.^16^ Mathematics was chosen for testing to reduce the effect of home learning on performance and better focus on classroom learning. Children were defined as having spectacles if they were able to produce them when asked, having previously been told to bring them to school that day.

### Visual acuity (VA) assessment

Children underwent VA screening by a local team of one nurse and one staff assistant who had previously been trained by optometrists and ophthalmologists from ZOC. Visual acuity was tested separately for each eye at a distance of 4 meters using Early Treatment Diabetic Retinopathy Study (ETDRS)^17^ tumbling E charts (Precision Vision, La Salle, Illinois) in a well-lighted, indoor area of each school. Visual acuity was measured without refractive correction for all children, and with habitually-worn correction for those children having eyeglasses.

Each child started testing from the 6/60 line. If the orientation of at least four of the five optotypes was correctly identified, the child was next examined on the 6/30 line. If one or no optotypes were missed, testing continued at 6/15 and proceeded line by line to 6/6. In case of failure to correctly identify 4 or more optotypes on a line, the line immediately above was tested until the child identified at least four of the five optotypes on a single line. The lowest line read successfully was recorded as the VA for the eye undergoing testing. If the top line could not be read correctly at a distance of 4 meters, the subject was directed to stand at 1 meter from the chart, and to read it as outlined above. In this case, the VA recorded was divided by 4.

Visual impairment (VI) was defined as presenting VA <= 6/12 in the better eye, and was stratified into mild (presenting VA <=6/12 to 6/18 in the better-seeing eye), moderate (<6/18 to 6/60) and severe (<6/60) according to categories proposed in the WHO International Statistical Classification of Disease 10^th^ Revision (ICD10).^18^

### Statistical Methods

We used log of the Minimum Angle of Resolution (logMAR) to denote VA. An increase of 0.1 logMAR units indicates a decrease of one line on an ETDRS chart. A higher logMAR value is indicative of worse vision.^19^

We compared various potential predictors of visual impairment in simple and multiple regression analyses. To explore the association between visual impairment and children’s scores on the standardized math test, we employed parametric Ordinary Least Squares regression methods. Regression models controlled for age, sex, spectacle ownership, boarding status, home county, parental education, and parental migration status. In order to explore potential factors determining ownership of spectacles, we employed Ordinary Least Square models including age, sex, uncorrected VA, boarding status, home county, parental education, parental migration status, including only students with uncorrected visual acuity <= 6/12 in the better-seeing eye that could be improved to >= 6/9 in at least one eye with glasses.

## Results

Among 2248 students undergoing vision screening (48% girls, mean age 13.6 +/- 1.1years), 100% completed vision screening, mathematics testing and all questionnaires. VI was present in 699 (31.1%; Table 1). Female sex (P < 0.001), older age (P = 0.05) home residence of Hengshan county, higher levels of maternal education (P = 0.006) and parental out-migration for work were associated with any VI, while boarding at school and paternal education level were not (Table 1).

**Table 1.** Characteristics of children with and without visual impairment (VI, presenting visual acuity <= 6/12 in both eyes)

| Characteristic | (1)<br>Normal<br>vision<br>N=1282<br>(57.0%) | (2)<br>Poor<br>uncorrected<br>vision<br>N=966<br>(43.0%) | (3)<br>visually<br>impaired<br>N=699<br>(31.1%) | (4)<br>Comparing<br>normal vision<br>with Poor<br>uncorrected<br>vision<br>P-value<br>(1)-(2) | (5)<br>Comparing<br>normal vision<br>with visually<br>impaired<br>P-value<br>(1)-(3) |
| --- | --- | --- | --- | --- | --- |
| **Children's Characteristics** |  |  |  |  |  |
| Age (years) | 13.514 | 13.636 | 13.634 | 0.01<sup>b</sup> | 0.03 |
| Male sex | 741 (57.8) | 432 (44.7) | 313 (44.8) | P<0.001 | P<0.001 |
| Boarding at school | 933 (72.3) | 718 (74.3) | 511 (73.1) | 0.41 | 0.88 |
| Home Hengshan county | 494 (38.5) | 248 (25.7) | 207 (29.6) | P<0.001 | P<0.001 |
| Home Jingbian county | 348 (27.2) | 289 (29.9) | 216 (30.9) | 0.15 | 0.08 |
| **Parental and Family Characteristics** |  |  |  |  |  |
| Mothers' education |  |  |  | 0.01 | 0.02 |
| Illiterate | 357 (28.2) | 219 (23.0) | 148 (21.5) |  |  |
| Not completing high school | 866 (67.6) | 693 (71.7) | 511 (73.1) |  |  |
| > 12 years of education | 43 (3.5) | 40 (4.1) | 28 (4.0) |  |  |
| Fathers' education |  |  |  | 0.51 | 0.23 |
| Illiterate | 110 (8.7) | 68 (7.1) | 44 (6.4) |  |  |
| Not completing high school | 1087 (84.8) | 813 (84.1) | 595 (85.2) |  |  |
| > 12 years of education | 71 (5.5) | 74 (7.7) | 53 (7.6) |  |  |
| Both parents out-migrated for work | 101 (7.9) | 92 (9.5) | 73 (10.4) | 0.17 | 0.05** |

Note: <sup>a</sup>\*Data are means (SD) or numbers (%) unless otherwise stated

Among children with poor uncorrected vision, 360 (37%) were observed owning spectacles. In simple regression models, uncorrected VA (P < 0.001), home residence in Hengshan county (P <0.0001), home residence in Jingbian county (P < 0.0001) were associated with wearing spectacles (Table 2). Age, male sex, boarding status, maternal education, paternal education, and parental migration status were un-associated with spectacle wear in the simple or multiple models (Table 2).

**Table 2.** Logistic regression model of potential predictors of spectacle wear among children needing glasses

|  | Simple analysis(n=966) |  | Multiple analysis (n=966) |  |
| --- | --- | --- | --- | --- |
|  | Regression coefficient (95%CI) | P-value | Regression coefficient (95%CI) | P-value |
| **Children's Characteristics** |  |  |  |  |
| logMAR | 0.958 (0.836,1.080) | P<0.001 | 4.893 (4.083,5.703) | p<0.001 |
| Age (years) | 0.005 (-0.021,0.030) | 0.73 | 0.001 (-0.132,0.133) | 0.99 |
| Male sex | -0.017 (-0.078,0.045) | 0.59 | -0.095 (-0.399,0.210) | 0.54 |
| Boarding at school | 0.084 (0.016,0.152) | 0.02 | 0.237 (-0.112,0.587) | 0.18 |
| Home Hengshan county | -0.181 (-0.246,-0.117) | p<0.001 | -1.335 (-1.730,-0.940) | p<0.001 |
| Home Jingbian county | -0.087 (-0.152,-0.022) | 0.01 | -0.811 (-1.170,-0.452) | p<0.001 |
| **Parental and Family Characteristics** |  |  |  |  |
| Mothers' education (Illiterate as reference) |  |  |  |  |
| Not completing high school | -0.016 (-0.085,0.051) | 0.63 | -0.238 (-0.610,0.133) | 0.21 |
| >12 years education | 0.002 (-0.151,0.156) | 0.98 | -0.013 (-0.839,0.812) | 0.97 |
| Fathers' education (Illiterate as reference) |  |  |  |  |
| Not completing high school | -0.015 (-0.099,0.069) | 0.72 | 0.060 (-0.587,0.707) | 0.86 |
| >12 years education | 0.006 (-0.109,0.121) | 0.92 | 0.245 (-0.575,1.065) | 0.56 |
| Both parents out-migrated for work | -0.076 (-0.175,0.024) | 0.14 | -0.397 (-0.985,0.192) | 0.19 |

Note: <sup>a</sup>logMAR is based on uncorrected VA in this table.

In multiple regression models of score on the mathematics examination, younger age (−0.227 SD, 95% Confidence Interval [CI] −0.283, −0.171, P < 0.001), wearing spectacles (0.147 SD, 95% CI 0.007, 0.288 P = 0.04), boarding at school (0.144 SD, 95% CI 0.003, 0.286 P = 0.05) and home residence in Hengshan county (0.217 SD, 95% CI 0.059, 0.374 P = 0.01) were associated with better scores, while paternal education, VA and sex were not (Table 3).

**Table 3.** Linear Regression Model of Predictors of Score on Mathematics Test (Expressed in units of Standard Deviation [SD])

|  | Simple Model n=966 |  | Multiple Model n=966 |  |
| --- | --- | --- | --- | --- |
|  | Coefficient<br>(95%CI) | P-value | Coefficient<br>(95%CI) | P-value |
| **Children's Characteristics** |  |  |  |  |
| Wearing spectacle | 0.145 (-0.050,0.229) | 0.03 | 0.147 (0.007,0.288) | 0.04 |
| logMAR | 0.218 (-0.076,0.511) | 0.15 | 0.144 (-0.161,0.448) | 0.36 |
| Age (years) | -0.221 (-0.277,-0.165) | p<0.001 | -0.227 (-0.283,-0.171) | P<0.001 |
| Male sex | 0.031 (-0.098,0.160) | 0.64 | 0.042 (-0.082,0.167) | 0.51 |
| Boarding at school | 0.117 (-0.027,0.260) | 0.11 | 0.144 (0.003,0.286) | 0.05 |
| Home Hengshan county | 0.154 (0.010,0.297) | 0.04 | 0.217 (0.059,0.374) | 0.01 |
| Home Jingbian county | -0.093 (-0.230,0.044) | 0.18 | 0.068 (-0.082,0.218) | 0.37 |
| **Parental and Family Characteristics** |  |  |  |  |
| Mother's education (Illiterate as reference) |  |  |  |  |
| Not completing high school | 0.024 (-0.118,0.165) | 0.74 | -0.019 (-0.161,0.122) | 0.79 |
| >12 years education | 0.089 (-0.297,0.475) | 0.65 | -0.033 (-0.445,0.379) | 0.88 |
| Father's education (Illiterate as reference) |  |  |  |  |
| Not completing high school | -0.078 (-0.263,0.107) | 0.41 | 0.026 (-0.209,0.261) | 0.83 |
| >12 years education | 0.265 (0.003,0.527) | 0.05 | 0.264 (-0.083,0.611) | 0.14 |
| Both parents out-migrated for work | -0.070 (-0.279,0.139) | 0.51 | -0.095 (-0.302,0.113) | 0.37 |

Note: <sup>a</sup>logMAR is based on presenting VA in this table.

## Discussion

Only about 37% of secondary school children needing glasses in this rural, western China cohort were wearing them, which is consistent with studies from other regions in China showing high rates of unmet need among rural children.^8,20^ In a study conducted in the same region, 15% of elementary school children needing glasses were observed wearing them.^8^ Unfortunately, data on the reasons for non-wear are not available from the current study, but a possible reason for the lower rates of spectacle use among younger children with refractive error is the well-documented and widespread belief in China that spectacle wear will harm the vision of young children.^3,21,22^ This view appears to be particularly strongly-held with respect to younger children, perhaps explaining the lower rates of wear among primary^8^ versus middle school children using the identical protocol in nearby areas of western China. A recent large trial has in fact shown spectacle wear to be protective of, rather than harmful to children’s vision,^23^ but the belief remains pervasive, and is focused principally on the youngest children. Our finding that wear of spectacles was highest in children with the most severe levels of uncorrected VA is also consistent with previous studies, though also as previously noted,^3,24^ rates of wear are from ideal.

These low rates of wear among visually impaired children is of particular concern in view of recent trial evidence that the provision of spectacles can significantly improve educational outcomes.^11,12^ Our finding in the current study that children owning spectacles had better performance on a study-specific mathematics test are consistent with this result, and with most,^13,25^ but not all^26^ of the limited previous non-trial data.

The difference in scores between children having uncorrected VA <= 6/12 in the better-seeing eye with and without spectacles in the current study (0.147 SD) was greater than the observed effect of parental education on math test score outcome, and is the equivalent to roughly a semester of additional learning.^27^ The significantly higher scores seen among younger children in each grade may reflect the fact that older children have been held back for poor school performance.

In the context of a cross-sectional, uncontrolled study such as the current investigation, we cannot exclude the possibility that the association between spectacle wear and learning could have been confounded by other factors, such as myopia or socio-economic status. Regarding the former, our data show that children with worse uncorrected VA (principally due to myopia in this setting) were more likely to wear spectacles, and myopia is also known to be associated with academic accomplishment.^28^ Similarly, higher socioeconomic status would be expected to be associated with greater likelihood of spectacle purchase, and has also been linked with academic accomplishment.^29^ Unfortunately, we did not collect data on either refractive error or socioeconomic status that might have allowed us to explore and adjust for such possible confounding.

These results have a clear message for program planners: much work remains to be done to improve wear of spectacles among rural Chinese children with refractive error, and the visual burden among those without spectacles is quite significant. Educational interventions directed at children, teachers and parents, and explaining the safety and value of spectacle wear have shown rather limited^5^ or no^30^ success in increasing rates of wear in recent randomized trials in China. Conversely, provision of free spectacles has recently been shown to double rates of both observed and self-reported wear in randomized trials.^11,12^ The educational benefits of spectacle wear in the classroom underscores the need for sustainable strategies to remove barriers to ownership and wear of spectacles.

Strengths of the current study include randomized population sampling, careful duplication of previously-utilized protocols and equipment^4^ in order to measure rates of wear for different age groups in a rural population, and our having administered a study-specific test in order to assess the impact of spectacle wear on education, as has been rarely done. The most obvious weakness of the current report is its cross-sectional design, making it difficult to elucidate with certainty the direction of any causal association between spectacle wear and educational outcomes. Further, we did not collect data on myopia and socio-economic status which might have allowed us to exclude these as sources of confounding in the observed relationship between spectacle wear and test outcomes. Finally, all of these schools were selected from a single prefecture in rural western China; hence application of these results to other settings must be made with caution.

Despite these limitations, results from this study provide further evidence of an association between spectacle wear and academic performance, as well as suggestive evidence of important differences in rates of spectacle wear between age groups in rural China. Both of these findings are of potential importance to program planners in formulating strategies to alleviate the burden of uncorrected myopia in rural China.

## Acknowledgments

We are grateful to all respondents who participated in this study and the enumerators for data collection efforts.

